# Role of CIA2 and CIL in the regulation of chloroplast photomorphogenesis in Arabidopsis

**DOI:** 10.1101/2025.11.24.690140

**Authors:** Roshanak Zarrin Ghalami, Pawel Burdiak, Muhammad Kamran, Maria Duszyn, Anna Rusaczonek, Ewa Muszyńska-Sadłowska, Stanisław Karpiński

## Abstract

Chloroplast development plays a crucial role in plant de-etiolation, a process in which plants switch from growth in darkness to light-driven development, known as photomorphogenesis. This study provides evidence that CIA2 (Chloroplast Import Apparatus 2) and CIL (CIA2-Like) contribute to chloroplast biogenesis, likely by affecting and regulating PSII assembly and related gene expression. Although their precise molecular roles remain unclear, our findings support their involvement in chloroplast development. This is indicated by deregulation of foliar chlorophyll content, chlorophyll *a* fluorescence parameters, chloroplast size, and gene expression of PSII molecular markers in *cia2cil* double mutant during de-etiolation. Chlorophyll *a* fluorescence and quantitative gene expression analysis during de-etiolation revealed a significant reduction in PSII maximal efficiency and non-photochemical quenching, as well as deregulated transcription of genes such as *LHCB2.1* and *psbA*. According to the immunoblotting and microscopy imaging results, there is an impaired assembly of PSII and a compromised ultrastructure of the chloroplast membranes in *cia2cil* plants. However, in *CIA2p::CIA2_cia2cil_* and *35Sp::CIA2_cia2cil_* complementation lines, reversion of this phenotype was observed. These results suggest a supporting role for CIA2 and CIL in the plant de-etiolation process, expanding our understanding of chloroplast biogenesis regulation.

## Introduction

Plants have evolved several ways of coping with different and highly fluctuating environmental conditions. The transition from a dark-grown, etiolated state to a light-adapted, photosynthetically active state, known as de-etiolation, is considered one of the most critical steps in plant development, enabling the plant to achieve its capability for photosynthesis. In this process, plants undergo some biochemical, physiological and morphological changes such as stem shortening, leaf expansion, chlorophyll synthesis, and functional chloroplast development [1–3]. De-etiolation involves optimizing a plant’s ability to receive and convert absorbed light into biochemistry and plant development. It is controlled by the complex cross-talk of several signaling pathways, including plastid and chloroplast retrograde signaling, as well as phytochromes and cryptochromes [1,2]. Regulatory mechanisms of chloroplast biogenesis are highly complex and include the interactions of numerous transcription factors (TFs) that regulate gene expression in response to light signals [2,4–6].

Chloroplast development is a process that requires the interaction of nuclear and plastid gene expression to provide the efficient assembly and optimal function of the photosynthetic apparatus [7,8]. Moreover, this process is also involved in transporting nuclear-encoded proteins across the double chloroplast membrane, mediated by specialized translocon complexes, to facilitate proper chloroplast biogenesis and function [9]. The translocons at the outer membrane of chloroplasts (TOC) and the inner membrane of chloroplasts (TIC) collaborate to import precursor proteins required for chloroplast formation and photosynthesis [10]. The TOC complex functions as a selective gateway, recognizing and transporting preproteins in a GTP-dependent manner, whereas the TIC complex controls their migration into the stroma [11]. Light triggers TOC159 activity to enhance the transport of photosynthetic proteins and hormonal signals like gibberellins and cytokinins regulate translocon function to balance chloroplast differentiation and development under changing environmental conditions [12,13]. The ultimate end of chloroplast development is regulated by the cell death mechanism that depends on cell death regulators (LSD1, EDS1, PAD4), non-photochemical quenching (NPQ) and PsbS [14–16].

Several TFs are involved in the chloroplast biogenesis process. HY5 is a transcription regulator of light-mediated genes, which is responsible for chlorophyll and photosynthetic proteins synthesis and thylakoid membrane formation [17–19]. Phytochrome-interacting transcription factors (PIFs) integrates light and hormonal signals, particularly gibberellin, to control chloroplast maturation and photomorphogenesis [20]. GLK1 and GLK2 (GOLDEN2-LIKE 1 and 2) enhance the expression of genes encoding chlorophyll biosynthesis enzymes and photosystem components [21–23]. CGA1 (cytokinin-responsive GATA factor 1) is a cytokinin-responsive transcription factor that upregulates photosynthesis-associated nuclear genes (PhANGs), along with MYB-related transcription factors (MYBS1), as regulators of chloroplast biogenesis, functioning in coordination with GLK transcription factors to ensure proper chloroplast biogenesis [24]. Other proteins like Early light-induced protein 1 (ELIP1), Photosystem II protein D1 (psbA), and LIGHT-HARVESTING CHLOROPHYLL A/B-BINDING (LHCB2) are also involved in protecting chloroplasts against photo-oxidative damage through efficient energy dissipation, and energy transfer within PSII for optimal photosynthetic activity [25–28]. Transcription factors, CIA2 and CIL, which are potentially dual-localized in the nucleus, have been suggested to regulate chloroplast protein import and biogenesis either directly or indirectly through GLK1 [29–31]. The phenotype of the *cia2* mutant exhibits a pale-green appearance, likely due to deregulated protein import, which may be attributed to impaired chloroplast development [30–32]. In addition to their roles in chloroplast biogenesis, CIA2 and CIL also participate in other aspects of plant growth and development, including the heat shock response and flowering [30,32]. However, the precise mechanisms by which CIA2 and CIL might regulate chloroplast biogenesis during de-etiolation remain unclear and require further investigation. Therefore, this study aims to investigate the roles of CIA2 and CIL in chloroplast development, focusing on their contribution to PSII assembly and chloroplast ultrastructure during de-etiolation.

## Results

### CIA2 and CIL promote cotyledon greening of etiolated seedlings during de-etiolation and are essential for chloroplast biogenesis

Previous studies indicate that CIA2 and CIL impact chloroplast development and plastid rRNA maturation, processes that may indirectly influence translational capacity [30]. We further investigated whether these proteins are involved in regulating chloroplast development during de-etiolation. The phenotypic analysis highlights the developmental dynamics of all genotypes used in this study (Figure 1A). All genotypes, except the *cia2cil* double mutant, showed progressive greening and an increase in chlorophyll and carotenoid content over time, consistent with active chloroplast biogenesis in response to light. The chlorophyll a/b ratio also gradually increased in all lines during de-etiolation, reflecting gradual chloroplast development. In contrast, the double mutant consistently exhibited lower chlorophyll a/b ratios, particularly between 24 and 96 hours (Figure 1B, S2). The *cia2cil* double mutant maintained a pale green phenotype in all analyzed time points compared to Col-0 (WT), whereas its phenotype was reverted in complementation lines. To determine the changes in CIA2 and CIL promoter activity during de-etiolation, a luciferase assay was performed using the CIA2 and CIL promoters. The analysis revealed that both these transcription factors are upregulated during the first hours of de-etiolation (T0-T8) (Figure 1C). This supports the involvement of CIA2 and CIL in the early stages of the photomorphogenic transition. To check the difference between complementation lines, we quantified *CIA2* transcript levels at T0-T96 time points in Col-0, 3*5Sp::CIA2_cia2cil,_*and *CIA2p::CIA2_cia2cil_*. *CIA2* expression in *35Sp::CIA2_cia2cil_*was highest across all time points except T0. While *CIA2p::CIA2_cia2cil_*showed lower expression overall, with a transient peak at 0 hours after light exposure (Figure S1).

**Figure 1:**
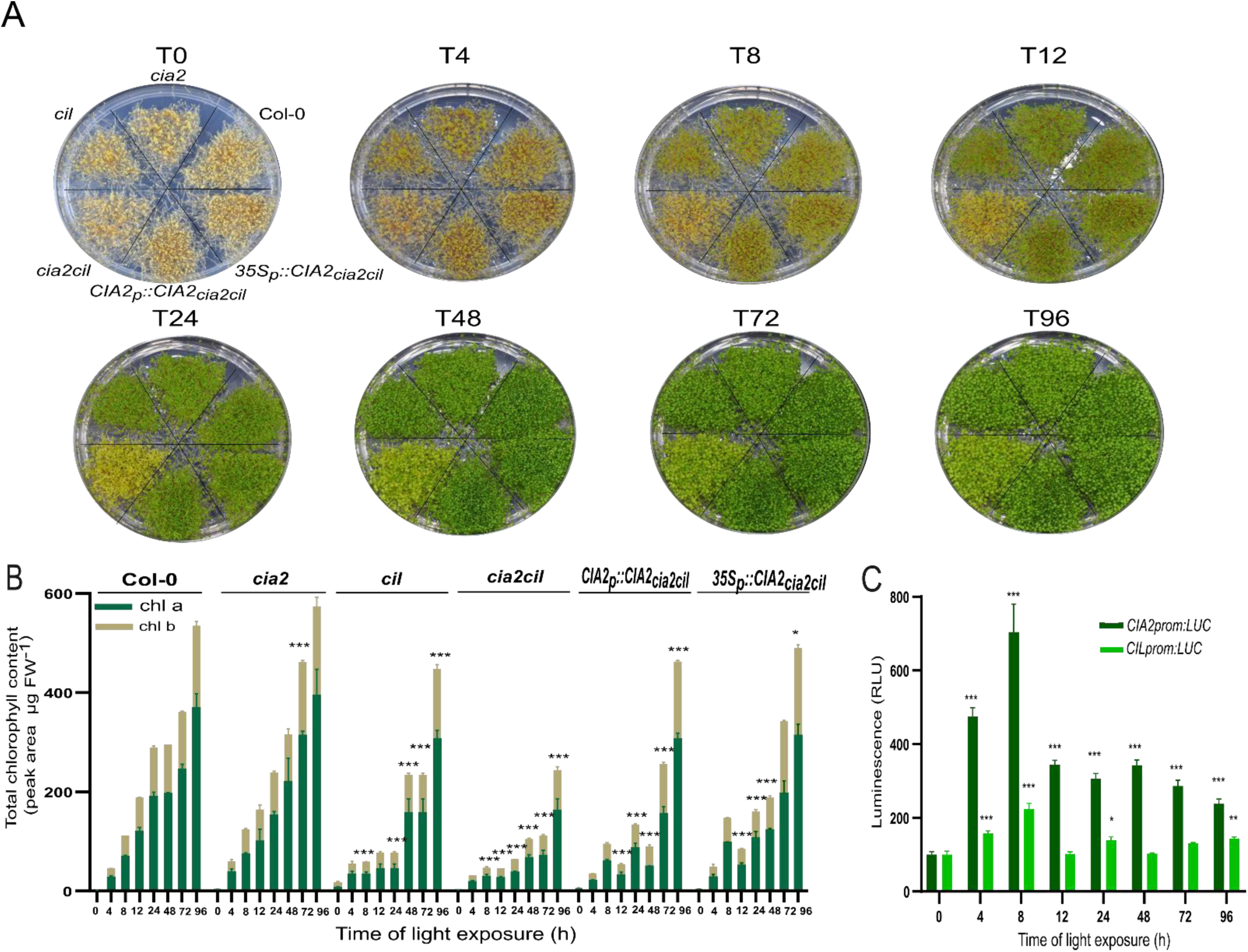
Pigment accumulation and CIA2/CIL promoter activity during de-etiolation in Arabidopsis seedlings. (A) *Arabidopsis thaliana* seedlings of the following genotypes: Col-0, *cia2, cil, cia2cil*, *CIA2p::CIA2_cia2cil_, 35Sp::CIA2_cia2cil_* during the first 96 hours of de-etiolation (T0, T4, T8, T12, T24, T48, T72 and T96). (B) Analysis of chlorophyll content. (C) Luciferase reporter assay showing *CIA2* and *CIL* promoter activity during the first 96 hours of de-etiolation. This study involved two genetic constructs carrying luciferase genes driven by *CIA2* or *CIL* promoters. Mean values were derived from 9 measurements (n = 9), and statistical significance (ANOVA and Tukey HSD test) is shown relative to Col-0 (P < 0.05 (*), P < 0.005 (**), or P < 0.001 (***)).

To investigate chloroplast development during de-etiolation in *cia2*, *cil,* and *cia2cil* mutants, confocal and transmission electron microscopy (TEM) were used. In confocal microscopy observations, chloroplasts in cia2cil exhibited abnormal structures and were noticeably smaller than those in the wild type (Col-0) and single mutants (Figure 2A-D, 2G). Complementation lines significantly reduced these structural defects, with *CIA2p::CIA2_cia2cil_* restoring chloroplast morphology and size to nearly wild-type levels (Figure 2A-G).

**Figure 2.**
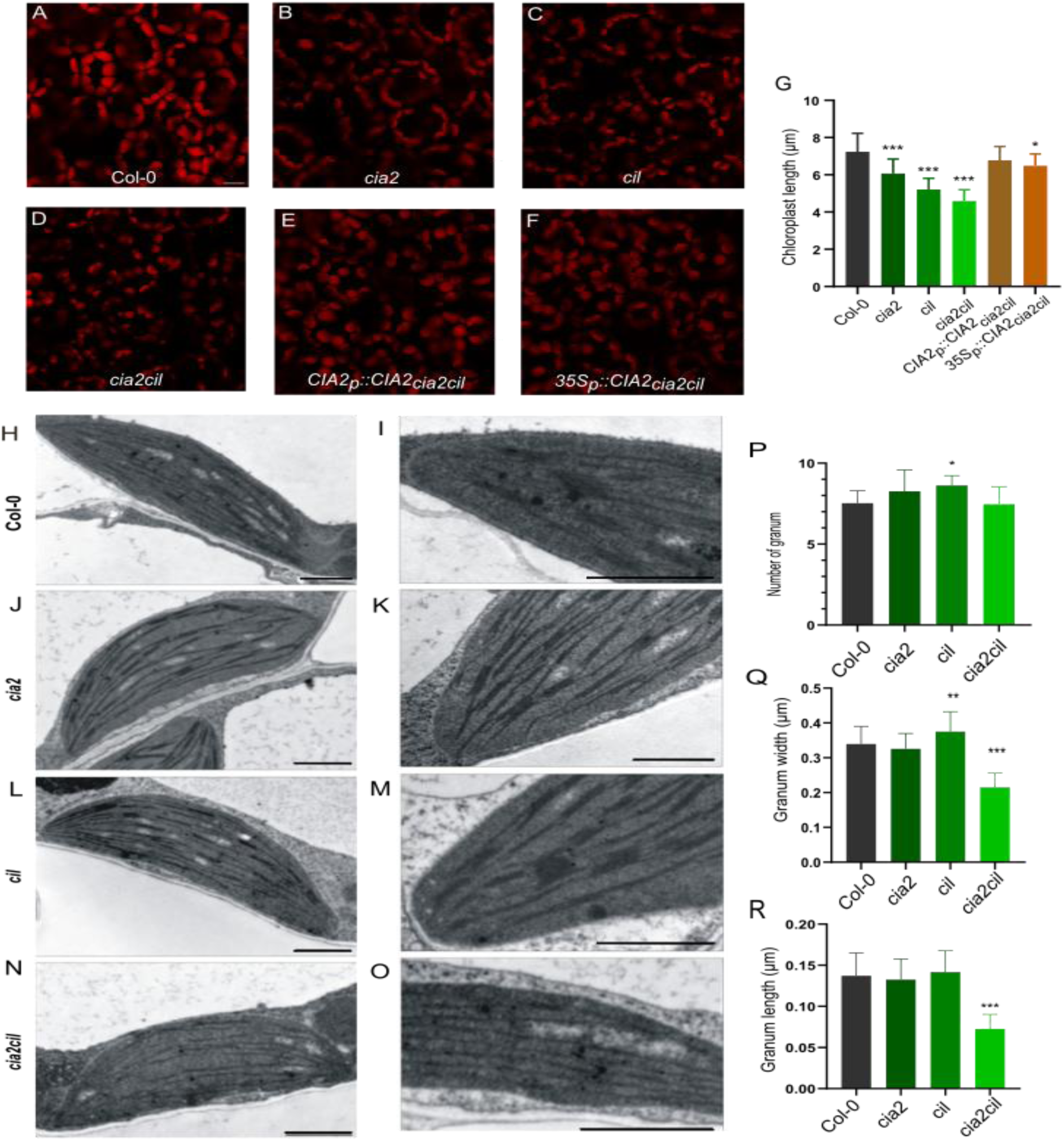
Ultrastructural analysis of chloroplast morphology during de-etiolation. (A-F) Confocal laser scanning microscopy (CLSM) images of chloroplasts (scale bar = 10 μm) in the (A) Col-0, (B) *cia2,* (C) *cil,* (D) *cia2cil* mutants and (E) *CIA2p::CIA2_cia2cil_* and (F) *35Sp::CIA2_cia2cil_ complementation* lines after 48h of de-etiolation. The red fluorescence corresponds to the chlorophyll fluorescence in the chloroplasts of mesophyll cells. (G) Quantitative analysis of chloroplast diameter (in μm), derived from confocal images using Fiji (ImageJ) (n=25). (H-O) Transmission Electron Microscopy (TEM) images of chloroplasts (scale bar = 1μm) in the (H,I) Col-0 and the (J,K) *cia2,* (L,M) *cil,* (N,O) *cia2cil* mutants after 48h. (P) The number of granum and quantitative analysis of granum (Q) length and (R) width (in μm), derived from TEM images using Fiji (ImageJ) (n = 40), and statistical significance (ANOVA and Tukey HSD test) is shown relative to Col-0 (P < 0.05 (*), P < 0.005 (**), or P < 0.001 (***)).

A detailed analysis of the thylakoid structure by TEM revealed that the length and width of the grana were significantly decreased in *cia2cil* compared to all other analyzed genotypes. However, the double mutant granum number (per chloroplast) remained unchanged, and interestingly, *cil* single mutants exhibited an increase in both grana width and length, while the *cia2* single mutant displayed no significant differences compared to wild-type (Figure 2 P-R). Moreover, TEM analysis of chloroplasts in the *cia2cil* showed defects in thylakoid membranes and less well-organized grana structures (Figure 2 N-Q, P-R).

### Role of CIA2 and CIL in maintaining PSII function during de-etiolation

To further confirm if CIA2 and CIL are required for optimal PSII function during de-etiolation, we monitored chlorophyll *a* fluorescence to visualize the dynamic responses of photosystems after transition from dark to light conditions. PSII maximal photochemical efficiency (*F*_v_/*F*_m_) increased gradually in wild-type plants and achieved a maximal level 24 h after light exposure, while *cia2cil* double mutant plants after 72 h (Figure 3A). In complementation lines, *Fv/Fm* values were fully restored; however, NPQ levels were only partially restored when compared to the wild type (Figure 3A, B). Additionally, the double mutant maintained increased non-photochemical quenching (NPQ) after 24h of light exposure (Figure 3B), indicating a prolonged reliance on protective energy dissipation processes. On the other hand, complementation lines, particularly *CIA2p::CIA2_cia2cil_*, exhibited a significant recovery in PSII efficiency, similar to that of wild-type plants.

**Figure 3.**
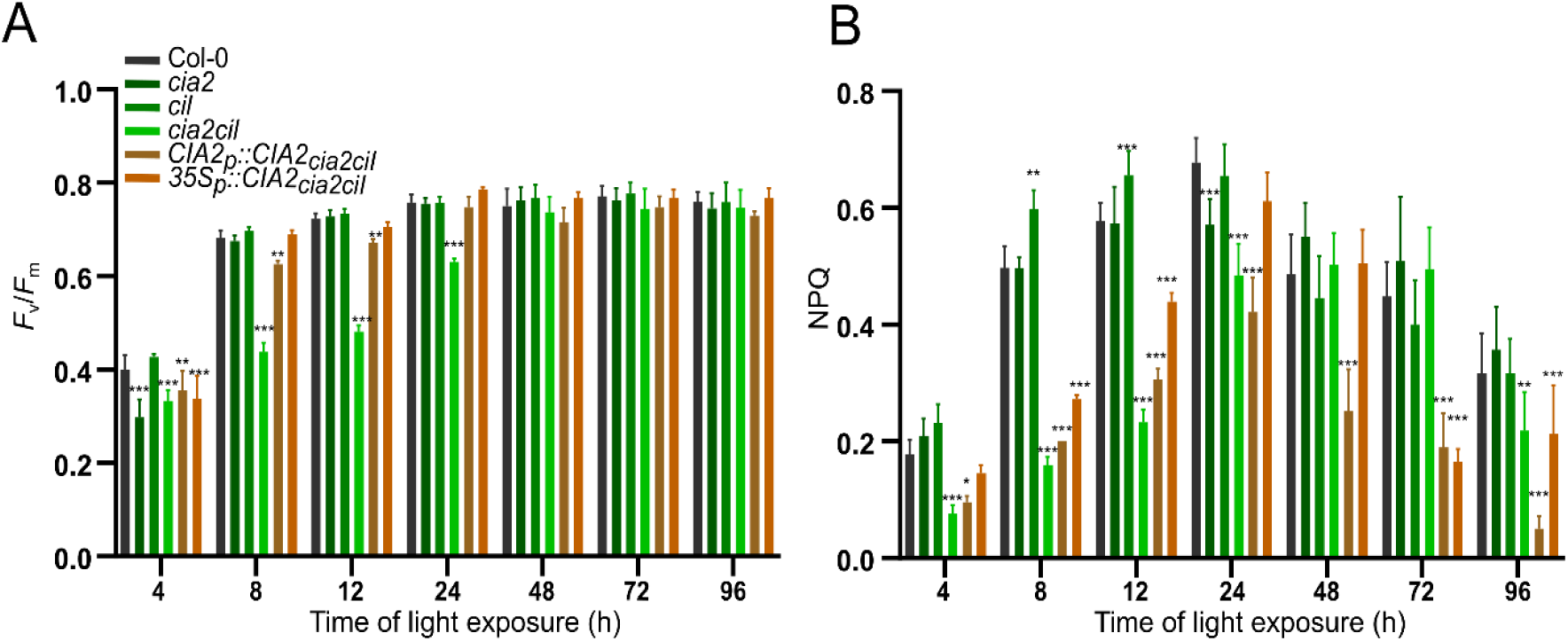
Photosystem II performance during de-etiolation monitored by chlorophyll a fluorescence parameters. (A) Maximum quantum efficiency of photosystem II (*F*_v_/*F*_m_) and (B) non-photochemical quenching (NPQ) in Col-0, *cia2*, *cil*, *cia2cil*, *CIA2p::CIA2_cia2ci_*_l,_ and *35Sp::CIA2_cia2cil_* seedlings during 96h of de-etiolation. Mean values were derived from 12 measurements (n = 12), and statistical significance (ANOVA and Tukey HSD test) is shown relative to the Col-0 (P < 0.05 (*), P < 0.005 (**), or P < 0.001 (***)).

### CIA2 and CIL might regulate the expression of photosynthesis-related genes during de-etiolation

CIA2 and CIL have been demonstrated to function as transcription factors that regulate the expression of genes encoding chloroplast-targeted proteins, including key components of the chloroplast translational machinery. They also exhibited dual subcellular localization to the nucleus and chloroplasts, suggesting that CIA2 and CIL might play a role in coordinating gene expression between the nucleus and chloroplasts, which is critical for chloroplast biogenesis and development [30,32,33]. RT-qPCRs were performed to investigate whether these proteins also regulate gene expression in response to de-etiolation. The study revealed severe abnormalities in gene expression levels, particularly in the double mutant plants (Figure 4, S3). During early time points (T0-T24), the *cia2cil* double mutant displayed significantly increased expression of *ELIP1*, which encodes early light-inducible protein that is involved in protecting PSII against photooxidative stress. On the other hand, transcriptomic analysis in *cia2cil* revealed a significant downregulation of genes, such as *GLK1* and *GLK2,* which are involved in chloroplast development*, as well as LHCB2* and *psbA*, which are necessary for the functional PSII (Figure 4, S3). Deregulated expression patterns observed in double mutant plants correlated with delayed greening and reduced chlorophyll accumulation. Two other genes, *TIC110* and *TOC159*, which encode the chloroplast protein import machinery system, were also downregulated in the double mutant, indicating a disorder in the import of nuclear-encoded proteins required for chloroplast development. Light signaling regulators, *HY5* and *PIFs,* showed the opposite expression pattern. In wild-type plants, *HY5* expression, which promotes photomorphogenesis and chloroplast biogenesis, was induced between T8 and T24 of de-etiolation, while *PIF1* and *PIF4*, *both negative regulators of photomorphogenesis*, were downregulated at these periods. This pattern was opposite in the *cia2cil* double mutant, indicating a failure to switch from skotomorphogenic (dark-adapted) to photomorphogenic (light-adapted) development. Furthermore, down-regulation of ribosomal genes such as *RPL11*, *RPL18*, *RPL28*, and *RPS6* in *cia2cil* plants directly correlates with the delayed chloroplast development and PSII assembly observed in the *cia2cil* mutant (Figure 4, S3). Notably, the expression of genes involved in chloroplast development, photosynthesis, and ribosome biogenesis was partially or fully restored in both complementation lines (*CIA2p::CIA2_cia2cil_*and *35S::CIA2_cia2cil_*), suggesting functional rescue by CIA2. In contrast, the expression of *PIF1* and *PIF4* remained downregulated in both lines, further supporting their negative regulatory relationship with CIA2 activity during de-etiolation (Figure 4, S3).

**Figure 4:**
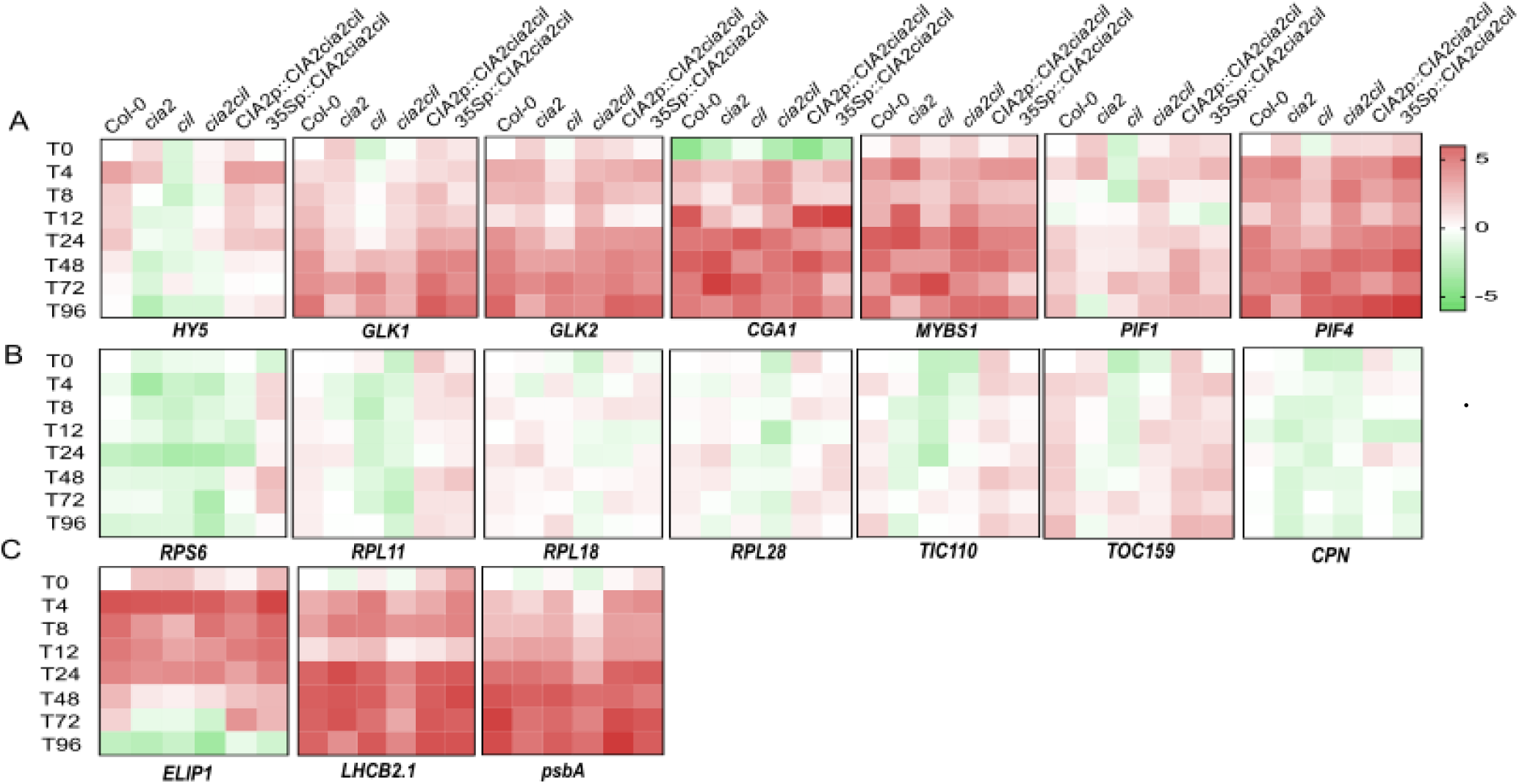
Gene expression heatmaps of chloroplast biogenesis-related pathways during de-etiolation. Heat-map of the expression of genes involved in (A) chloroplast development and transcriptional regulation, (B) protein translation and import, and (C) photosystem components and light harvesting in *cia2*, *cil*, *cia2cil*, *CIA2p::CIA2_cia2cil_, 35Sp::CIA2_cia2cil_* in 0-96 hours.

During de-etiolation, CGA1 expression showed strong suppression in the *cia2*, *cil*, and especially *cia2cil* mutants at 12 hours, a key time point for light-induced chloroplast gene activation. The complementation line *CIA2p::CIA2_cia2cil_* restored CGA1 expression to wild-type levels, while *35S::CIA2_cia2cil_* showed variable expression, particularly at later time points.

MYBS1 expression was elevated in *cia2* seedlings at the first time points (0-12 h), but was generally reduced in the *cil* mutant and unstable in *cia2cil*, indicating disrupted regulation. The *CIA2p::CIA2_cia2cil_*line maintained expression close to wild-type across the time course, while *35S::CIA2_cia2cil_* exhibited inconsistent and often lower expression.

Immunoblot analysis (Figure 5A) revealed a time-dependent pattern of ELIP1 protein accumulation, with high levels during the early hours of de-etiolation, followed by a gradual decline in all genotypes. The cia2cil double mutant consistently exhibited higher ELIP1 levels than the other genotypes throughout the time course. The accumulation of LHCB2 and psbA proteins in the double mutant was delayed, and protein buildup could be seen only after 72 hours of light exposure. Single mutants have a level of LHCB2 and psbA protein that is nearly identical to that of Col-0 throughout the de-etiolation process. In complementation lines, LHCB2 and psbA began to accumulate within 12 hours of light exposure and continued to increase with time, similar to Col-0.

**Figure 5.**
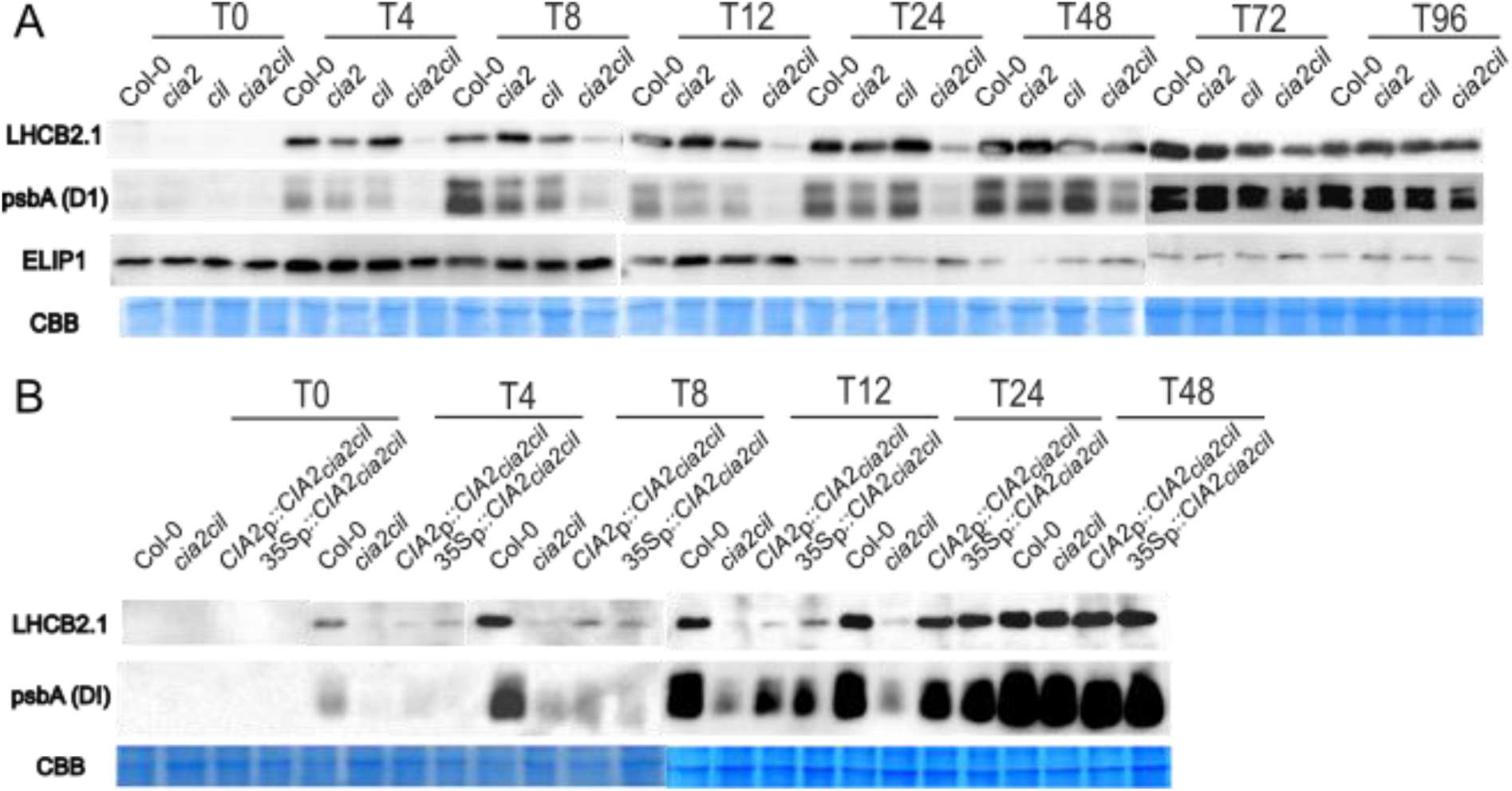
Immunoblot analysis of chloroplast protein accumulation during de-etiolation. Immunoblot analysis using total protein isolated from seedlings of (A) Col-0, *cia2*, *cil*, *cia2cil* during 96h of de-etiolation and (B) complementation lines (Col-0, *cia2cil*, *CIA2p::CIA2_cia2cil_, 35Sp::CIA2c_ia2cil_*) in T0-T48 time points. The level of ELIP1, LHCB2 and the D1 protein of PSII reaction center are shown. The level of the 50 kDa unknown protein, stained with Coomassie blue, confirmed equal gel loading.

## Discussion

The study reveals the potential roles of CIA2 and CIL in chloroplast biogenesis. Our data show that CIA2 and CIL function synergistically in chlorophyll synthesis, chloroplast development, and PSII activity. The *cia2cil* double mutant displays a pale-green phenotype, reduced pigment levels, disrupted chloroplast structure, and decreased PSII efficiency. It’s consistently lower chlorophyll a/b ratio compared to other genotypes suggests impaired PSII core assembly and an imbalance between PSII and its associated light-harvesting complexes. Complementation of the double mutant restored chloroplast structure, pigment content, and PSII performance to near wild-type levels. These findings suggest that CIA2 and CIL may play roles in PSII biogenesis and chloroplast maturation, either directly or indirectly (Figure 1-3). Furthermore, luciferase reporter assays revealed that the CIA2 and CIL promoters are strongly induced shortly after the transition from dark to light (Figure 1C). These findings support a role for CIA2 and CIL in early light-driven chloroplast development during de-etiolation.

These findings offer new insights into the roles of CIA2 and CIL in photomorphogenesis, extending previous research that highlighted their importance in chloroplast biogenesis only in mature plants [29,29,30,33]. Numerous regulators have been described to play roles in chloroplast biogenesis. Among these, GLKs synergistically influence the balance of chlorophyll biosynthesis, chloroplast development, and the level of PhANGs (Photosynthesis Associated Nuclear Genes) [23]. Furthermore, it has been suggested that GLKs play a more direct role in coordinating and regulating the expression of genes crucial for chloroplast development, chlorophyll biosynthesis, light harvesting, and carbon fixation (*GUN4, CAO, LHCB1-6*) [22]. Yang et al. (2022) suggested that CIA2 and CIL might regulate chloroplast development directly or indirectly *via* GLK1. Furthermore, their results implied that CIA2 plays a more crucial role than CIL in upregulating *GLK1* expression [22,33]. Disturbed chlorophyll synthesis, along with lower transcript and protein levels associated with light-harvesting complex (LHCB2) and photosystem assembly (D1) in *cia2cil* (Figure 4, 5), led to delayed and diminished PSII efficiency during de-etiolation (Figure 3), indicating a disturbed capacity of energy utilization and light capture [34,35]. A failure to assemble fully functioning PSII complexes in mutant plants results in elevated non-photochemical quenching (NPQ) after 24 h of exposure, indicating a dependence on protective energy dissipation processes. These results are consistent with those of Wang et al. (2022), which also demonstrated PSII inefficiency in mutants with chlorophyll deficiencies and highlighted the dependence of PSII functionality on pigment and protein biosynthesis pathways [36]. Lutein plays a role in stabilizing the LHCII and in enabling thermal energy dissipation in protecting photosystem II under high-light conditions [37–39]. Lower NPQ in *cia2cil* plants was accompanied by lower lutein contents (Figure 3B, S2), and lower LHCB2 level (Figure 5A), when compared to Col-0, suggesting impaired photoprotective mechanisms. Therefore, a lower level of LHCB2, which is required for the proper organization of antenna complexes, presumably resulted in impaired PSII assembly, as evidenced by the visible stacked thylakoid grana [34]. This association implies that the *cia2cil* photoprotection is compromised due to deficiencies in both carotenoid production and LHCII integrity. Additionally, decreased Fv/Fm, an indicator of PSII dysfunction, is commonly correlated with lower D1 protein levels [40–42]. In our studies, the *cia2cil* showed delayed development of PSII efficiency, along with a gradual increase of D1 accumulation from T0 to T96 (Figure 5A), which suggests that during this process, the CIA2 and CIL play a positive and putative regulatory role in D1 synthesis. On the other hand, the incomplete rescue of NPQ with complementation lines suggests that CIA2 alone is insufficient to fully restore the photoprotective capacity of the double mutant. NPQ involves rapid energy dissipation mechanisms that may rely more on CIL.

According to immunoblot analysis (Figure 5), the delayed accumulation of LHCB2 and psbA in the *cia2cil* double mutant suggests a deficiency in the nuclear and plastid protein translation during chloroplast development. Single mutants did not exhibit this delay, indicating that CIA2 and CIL have overlapping activities. Complementation with CIA2 partially improved early protein accumulation, especially after 12 hours of de-etiolation, which suggests that CIA2 might has a function in regulating photosynthetic protein synthesis since it has been shown that the translation of chloroplast mRNA and the maturation and accumulation of 23S rRNA are influenced by CIA2 and CIL [30]. Overall, CIA2 is critical for maintaining a balance between photoprotective and photosynthetic protein synthesis throughout the transition from etiolation to photomorphogenesis.

Confocal microscopy revealed abnormally formed chloroplasts in the *cia2cil* mutant, with decreased development and overall size (Figure 2). A smaller granum size only in *cia2cil* double mutant but not in single mutants implies that CIA2 and CIL regulatory proteins might have complementary functions in regulating the stacking and organization of thylakoid membranes and granum structure; however, they are not necessary for granum initiation, as shown by the unchanged granum number (Figure 2P). The single mutants, *cia2* and *cil*, exhibit fewer structural defects and show better photosynthetic function, suggesting that both CIA2 and CIL contribute to chloroplast development, and their combined loss exacerbates the phenotype. Ultrastructural observation using TEM revealed significant abnormalities, including undeveloped thylakoid membranes and diminished stacking at T48 (Figure 2P-R), consistent with another study that suggested similar defects in fully developed plants [31]. These findings are in accordance with reduced PSII assembly (Figure 3A) and emphasize the possible regulatory impact of CIA2 and CIL on chloroplast structure and function. A high level of ELIP1 in the double mutant (Figures 4 and 5A) suggests that plants attempt to reduce photooxidative damage, which is consistent with the findings of Kleine et al. (2007), who found comparable responses in chloroplast-stressed mutants. In both complementation lines, ELIP1 expression is partially or fully rescued, indicating that CIA2 alone can restore ELIP1 expression and likely its downstream chloroplast-related functions, even in the absence of CIL, thereby reinforcing the idea that CIA2 is a possible key regulator.

Beyond defects in chlorophyll synthesis and photosystem assembly, our data suggest broader disruptions in the chloroplast biogenesis machinery. In particular, the observed downregulation of key components of the protein import apparatus, such as *TIC110* and *TOC159* in all analyzed mutants (Figure 4, S3) is consistent with Sun et al. (2009) study, which showed CIA2 crucial role in protein import and translation processes. Also, the lack of expression of ribosomal genes such as *RPL11, RPL18, RPL28*, and *RPS6* reveals another reason for chloroplast development deficiency and delay in the *cia2cil* mutant, which is associated with inefficient chloroplast-encoded protein synthesis and assembly. These findings are consistent with Gawronski et al. (2021) study, which found that ribosomal deficits in *cia2cil* lead to a delay in chloroplast-targeted protein synthesis and assembly. CPN also showed decreased expression in *cia2cil* which is associated with the failure of post-import protein folding and maturation, a side effect of deficient protein import machinery [43].

The combined disruptions in protein import, folding, and assembly suggest that chloroplast development in *cia2cil* may be impaired not only locally within the organelle but also through defects in nuclear regulatory programs. The phenotypic similarities between *cia2cil* and *glks* mutants also support the hypothesis that CIA2 and CIL are involved in chloroplast biogenesis. Both double mutant lines exhibit pale green phenotypes and deficiencies in the formation of the photosynthetic apparatus [22,32]. Furthermore, their putative localization to chloroplasts implies possible functions as environmental sensors, directing adaptive responses via retrograde signaling [44]. Fitter et al. (2017) showed that these GLKs are required for chloroplast formation and the activation of photosynthetic machinery genes, while Yang et al. (2022) found that CIA2 and CIL coordinate chloroplast biogenesis and function primarily by enhancing the expression of the nuclear factor *GLK1* and genes associated with chloroplast transcription, translation, protein import, and photosynthesis.

Downregulation of CGA1 in *cia2cil* mutants emphasises the involvement of CIA2 and CIL in activating early light-responsive genes during de-etiolation, supporting their function in transcriptional coordination [45]. MYBS1 expression is upregulated in the *cia2* single mutant but is downregulated in the *cia2cil* double mutant, indicating a compensatory response to the loss of CIA2 alone, and a loss of regulatory control when both CIA2 and CIL are absent, which is consistent with MYBS1 functioning along with GLKs in Arabidopsis [24]. The reduced expression of GLKs in the double mutant reinforces the upstream role of CIA2 and CIL, while *CGA1*, *MYBS1*, *GLK1* and *GLK*2 expression were restored in the *CIA2p::CIA2_cia2cil_*complementation line but not in *35S::CIA2_cia2cil_*, emphasizing the importance of native promoter-driven expression for proper timing and coordination of chloroplast biogenesis. The differences between *CIA2p::CIA2_cia2cil_* and *35S::CIA2_cia2cil_*lines likely reflect that constitutive overexpression of CIA2 may disrupt regulatory balance, either by overriding normal developmental controls or by triggering negative feedback that suppresses downstream gene activation. Together, these results reveal that CIA2 and CIL may redundantly regulate key transcription factors (CGA1, MYBS1, GLK1 and GLK2), and that their absence disrupts chloroplast development by impairing the transcriptional network required for de-etiolation. Both complementation lines restored *HY5* activation and photomorphogenic development, but the *35Sp::CIA2_cia2cil_* line showed stronger upregulation of *PIF4*, likely reflecting excessive *CIA2* expression. This suggests that while proper *HY5* activation is sufficient for recovery, native levels of *CIA2* provide a more balanced regulation of the transcriptional network.

Beyond GLK-mediated regulation, the transcriptional balance between HY5 and PIFs is another critical component of light-dependent development. HY5 promotes light responses, whereas PIFs drive skotomorphogenesis; their opposite expression patterns in *cia2cil*, with *HY5* downregulated and *PIFs* upregulated in the double mutant, indicate impaired photomorphogenic development and an important role of CIA2 and CIL in balancing these opposing signaling pathways.

These transcriptional patterns indicate a coordinated regulatory network in which CIA2 and CIL might play a role. The downregulation of *TIC110, TOC159,* ribosome-encoded genes, and *CPN*, which leads to impaired protein synthesis and import, along with *GLK1, GLK2*, and *HY5* low expression and the upregulation of *PIFs* in the double mutant, indicates a widespread failure in chloroplast biogenesis. While in complementation lines, partial restoration was observed and highlights the importance of CIA2 in chloroplast protein import (e.g., *TOC33* and *TOC75*), plastid translation, protein folding, and photosynthesis functionality and defective ribosome assembly [30–32].

The observed differences in gene expression between the *CIA2p::CIA2_cia2cil_*and *35Sp::CIA2_cia2cil_* complementation lines likely reflect variation in promoter strength and positional effects. The greening phenotype observed in both lines indicates that a threshold level of *CIA2* expression is sufficient to support chloroplast development. However, the elevated and constitutive expression driven by the 35S promoter may alter the transcriptional fine-tuning required during de-etiolation, thereby contributing to the observed transcriptional differences. These interpretations are supported by the *CIA2* expression levels shown in Figure S1. The gene expression patterns of complementation lines highlight the role of CIA2 as an important positive and putative regulator of chloroplast biogenesis and photomorphogenesis. In both *CIA2p::CIA2_cia2cil_* and *35S::CIA2_cia2cil_*lines, CIA2 complementation led to increased expression of genes related to photosynthesis (*GLK1*, *GLK2*, *LHCB2*, *psbA*), ribosome biogenesis (*RPS6*, *RPL11*, *RPL18, RPL28*), and protein import into the chloroplast machinery (*TIC110*, *TOC159*). Luciferase activity results show that *CIA2* is much more upregulated than *CIL* during the de-etiolation process (Figure 1C). This corresponds with the CIA2 putative role in the activation of genes required for chloroplast protein import and ribosomal protein production, and CIL has more supplementary role in this process [29,31–33]. In conclusion, our results show that CIA2 and CIL either directly or indirectly affect several aspects of chloroplast gene expression and PSII assembly (Figure 6). These TFs may have multiple roles, but their precise molecular functions and regulatory mechanisms require further investigation. Nonetheless, our study highlights the role of CIA2 and CIL in chloroplast biogenesis through the coordinated regulation of key genes. By integrating phenotypic, biochemical, molecular, and structural data, we provide new insights into the mechanisms of chloroplast formation. These findings also open up potential avenues for improving photosynthetic efficiency and stress tolerance in plants through genetic and biotechnological approaches.

**Figure 6.**
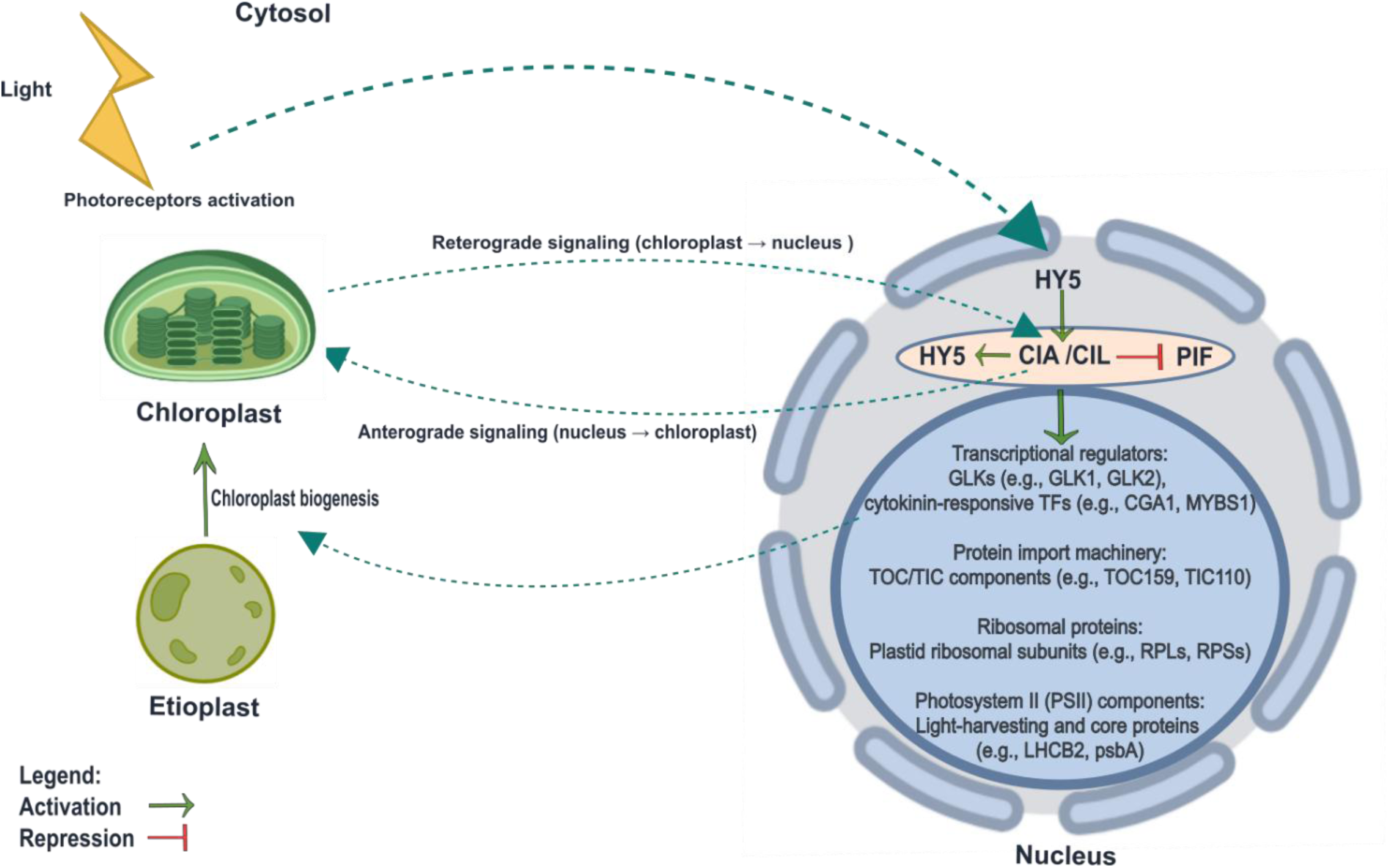
Proposed model of CIA2 and CIL regulatory roles in chloroplast biogenesis during de-etiolation in Arabidopsis thaliana. Light perception activates photoreceptors, leading to induction of HY5, a key transcription factor that promotes the expression of CIA2 and CIL. In turn, CIA2 and CIL reinforce HY5 expression and inhibit PIFs, negative regulators of photomorphogenesis, thereby promoting a successful transition from skotomorphogenesis to photomorphogenesis. CIA2 and CIL also activate nuclear genes required for chloroplast development, including GLK1/2, CGA1, MYBS1 (transcriptional regulators), TOC159/TIC110 (protein import machinery), RPLs/RPSs (ribosomal proteins), and LHCB2/psbA (PSII components). These regulatory cascades coordinate chloroplast biogenesis and etioplast-to-chloroplast transition. Bidirectional communication between the nucleus and chloroplast via anterograde and retrograde signaling ensures tight coordination of gene expression and organelle development.

## Material and Methods

### Plant material and growth conditions

*Arabidopsis thaliana* seeds (Columbia-0 (Col-0)*, cia2* (SALK_004037), *cil* (SAIL_228_C01)*, cia2cil, CIA2p::CIA2_cia2cil_,* and *35Sp::CIA2_cia2cil_* were surface-sterilized with chlorine gas generated by the reaction of sodium hypochlorite and concentrated hydrochloric acid (HCl). Seeds were sown in spots containing 50 seeds (to facilitate rapid harvest) on agar plates containing 0.5 Murashige and Skoog salt mixture (Duchefa Biochemie, Haarlem, Netherlands) without sucrose. Following stratification in the dark for 3 days at 4°C, plates with seeds were kept for 2 hours at the light intensity of 40 µmol photons m^-2^ s^-1^ at 21°C and then transferred to the dark for 3 days at 21°C. After 3 days of darkness, plates were transferred to long photoperiod (16/8 h) growing conditions with the light intensity of 80 µmol photons m^-2^ s^-1^. Seedlings were collected at each time point (T0, T4, T8, T12, T24, T48, T72, T96) in three replicates and transferred into a 1.5 ml tube, flash-frozen in liquid nitrogen, and stored at -80°C until further analysis [3,46].

### Construct Generation and Transformation

To generate the *CIA2p::CIA2_cia2cil_* and *35S::CIA2_cia2cil_*constructs, we employed the Golden Gate cloning method for precise assembly. Genomic DNA was extracted from *Arabidopsis thaliana* Col-0 using the CTAB method. We then amplified the full genomic sequence of *CIA2* along with its native promoter region (1670 bp upstream of the start codon) by PCR For the *CIA2p::CIA2_cia2cil_* construct, the native promoter was cloned upstream of the genomic *CIA2* sequence, preserving its natural regulatory control, whereas for the *35S::CIA2_cia2cil_*construct, it was replaced with the constitutive CaMV 35S promoter. Both constructs were assembled in a single Golden Gate reaction using the Type IIS restriction enzyme BsaI and the binary vector pGoldenGate-SE9. Afterward, the constructs were introduced into *cia2cil* double mutant background using *Agrobacterium tumefaciens* strain GV3101. Further experiments were conducted using the T3 generation of plants. For luciferase studies, the 2701 bp promoter region of *CIA2* and the 2566 bp promoter region of CIL were amplified with primers listed in Table S1 and S2. PCR products were purified and inserted into the entry clone using the pENTR/D-TOPO cloning kit (Invitrogen). Next, *CIA2* and *CIL* promoters were subcloned into the pGWB635 vector, with the luciferase reporter gene located upstream. The transgenic lines *CIA2p::LUC* and *CILp::LUC* were generated by floral dip transformation using Agrobacterium tumefaciens (GV3101). [47–49].

### Luciferase Activity Assay

To analyze luciferase activity, the Promega Luciferase Assay System kit was used. Seedlings were collected at various time-points (T0, T4, T8, T12, T24, T48, T72, T96). 35 mg of seedlings were homogenized in liquid nitrogen, followed by the addition of 250 µL cell lysis buffer. The supernatant was separated after centrifugation and mixed with a luciferin substrate solution. Luminescence was quantified using a luminometer (Berthold Technologies, Lumat LB9507) [50].

### Chlorophyll *a* fluorescence

Chlorophyll a fluorescence parameters were measured on seedlings using a pulse amplitude-modulated FluorCam 800 MF and the associated software (Photon Systems Instruments, Drasov, Czech Republic). Before measurements, the plates were kept in the dark for 30 min to determine *F*_0_ and *F*_m_. Chlorophyll fluorescence terminology has been previously described [51,52].

### Pigments analysis

20 –50 mg of frozen tissue was homogenized in a Mixer Mill MM 400 (Retsch, Düsseldorf, Germany) (5 min, 4 °C, 30 Hz) with 1 ml of cold acetone (−20 °C). The homogenate was evaporated using Savant DNA120 SpeedVac (Thermo Scientific, Waltham, MA, USA), dissolved in cold solvent A (acetonitrile: methanol; 90:10; v/v), and re-homogenized for 1 min. The extract was filtered through a 0.2 μm nylon filter (Whatman) into an auto-sampler vial, capped, and stored in the dark at −80 °C for HPLC analysis (Shimadzu, Kyoto, Japan). The pigments were separated on a Synergi^TM^ 4 μm MAX-RP 80 Å LC Column 250 × 4.6 mm (Phenomenex, Torrance, CA, USA) at 30 °C. Solvent A was used for 10 min to elute all xanthophylls, followed by solvent B (methanol: ethyl acetate; 68:32; v/v) for 10 min at a flow rate of 1 ml/min. The results are given as the peak area per μg of fresh weight, according to the protocol previously used [53,54].

### Immunoblot analysis

Proteins were extracted from whole seedlings in four volumes (w/v) of SDS-PAGE sample buffer (0.2M Tris/HCL pH 6.8, 0.4 M dithiothreitol, 8% (w/v) SDS, 0.4% (w/v) Bromophenol blue, and 40% (v/v) glycerol). Proteins were denatured for 15 min at 95°C, and cell debris was removed by centrifugation for 5 min at 16,000 g. Proteins (40 µg) were separated on SDS-PAGE (10–15% (w/v) polyacrylamide concentrations depending on the molecular weight of the protein of interest and transferred onto an Immobilon-P PVDF membrane (Merck) by semi-dry transfer. Immunodetections were performed using specific antibodies and dilutions: 1:500 LHCB2 (AS01 003, Agrisera), 1:10000 D1(psbA) (AS05 084, Agrisera) and 1:1000 ELIP1 (PHY0842A, PhytoAB). After incubation with primary antibodies overnight at 4°C, membranes were incubated for 1 h at RT with horseradish peroxidase–conjugated secondary antibodies (1:3000 (v/v) anti-rabbit secondary antibodies, Agrisera). Protein bands were immunodetected using SuperSignal West Dura Extended Duration Substrate (Thermo Scientific) according to the manufacturer’s recommendations, visualized with the ChemiDoc XRS+ System (Bio-Rad) and analyzed with ImageLab Software 5.2.1 (Bio-Rad)[55,56].

### Quantitative Real-Time PCR

RNA extraction was performed using Plant RNA Reagent (Life Technologies) according to the manufacturer’s instructions. cDNA was synthesized using a High-Capacity cDNA Reverse Transcription Kit (Life Technologies). All PCR amplifications were run on a 7500 Fast Real-Time PCR System (Applied Biosystems) using a Power SYBR Green PCR master mix (Life Technologies) UPL7 and PP2AA3 were used as reference genes to calculate relative expression [57]. All primers used in this study are listed in Table S1.

### Transmission electron microscopy

Sample preparation for TEM analysis involved fixation of leaf fragments (seedlings (T48)) in 2 % (v/v) glutaraldehyde and 2 % (v/v) paraformaldehyde in 0.1 M cacodylate buffer (pH 7.2) for 3 h and washing off four times in the same cacodylate buffer [58]. Next, the plant samples were secondary fixed in 2 % (v/v) osmium tetroxide for 2 h at low temperature, dehydrated in an ethanol gradient, and replaced with propylene oxide. Finally, samples were embedded in EPON epoxy resin. Its polymerization was performed at 60 _°_C overnight. Ultra-thin sections of about 80 nm thickness were prepared using a UCT ultramicrotome (Leica Microsystems), stained with uranyl acetate dissolved in a saturated ethanol solution, followed by incubation in lead citrate. An FEI 268D ‘Morgagni’ transmission electron microscope (FEI Company, Hillsboro, OR, USA) equipped with an Olympus-SIS ‘Morada’ digital camera (Olympus) was applied for the ultrastructural examination [58]. Quantitative analysis of thylakoids was performed using Fiji (ImageJ) [59].

### Confocal microscopy

Seedlings (T48) were analysed for chloroplast structure. Confocal microscopy observations were performed using a Zeiss LSM700 microscope equipped with 20x and 40x EC Plan-Neofluar objectives. Chlorophyll fluorescence was excited using a 488 nm laser. Signals were detected using 652-682BP (chlorophyll) filters with a beam splitter set at 601 nm. Quantitative analysis of chloroplast diameter (in μm), derived from confocal images, was performed using Fiji (ImageJ) [59,60].

### Statistical analysis

All statistical analyses were carried out using GraphPad Prism version 8. ANOVA and Tukey’s HSD test were used to examine differences across several experimental groups, followed by Tukey’s post-hoc test for pairwise comparisons. The data is provided as mean ± SEM, with statistical significance set at (P < 0.05 (*), P < 0.005 (**), or P < 0.001 (***)).

## Supplementary data

Table S1 and S2: Primer sequence used for quantitative real-time PCR (qRT-PCR) and constructs. Fig. S1. CIA2 expression in complementation lines. Fig. S2. Pigments analysis, the content of carotenoids and lutein. Fig. S3. Transcriptome Analysis. qRT-PCR analysis of gene expression in graphs.

## Author contributions

RZ, MK, MD, PB, AR, and EMS: investigation; RZ, PB, and MD: formal analysis and visualization; RZ: writing the original draft; PB, MD, and SK: writing - review & editing; SK: supervision and funding acquisition.

## Conflict of interest

The authors declare no conflict of interest.

## Funding

This work was supported by the Polish National Science Center (Narodowe Centrum Nauki; OPUS20, UMO-2020/39/B/NZ3/02103) given to S.K.

## Notes

### Competing Interest Statement

The authors have declared no competing interest.

