## Supplementary file for "Role of CIA2 and CIL in the regulation of chloroplast photomorphogenesis in Arabidopsis"

**Supplementary data:**

Table S1: List of primers used for quantitative real-time PCR (qRT-PCR) analysis of selected *Arabidopsis thaliana* genes.

| Gene | Accession Number | Forward Primer (5'→3') | Reverse Primer (5'→3') |
| --- | --- | --- | --- |
| *ELIP1* | AT3G22840 | GGCTATGCTGGGATTCGTGT | GACGCCAGGAACCAGATCAA |
| *LHCB2.1* | AT2G05070 | GGCTTTCTCTGAGCTGAAGG | GCGTTGTTAGCCACAGGATC |
| *PsbA* | ATCG00020 | GGTGCTCATTTTGTATGGGC | GGTTACTCCTACAGCACGTC |
| *RPL11* | AT1G32990 | CCTCAACTCCGAGATTCTCAC | AAGCAGTTTGATAATCCACA |
| *RPL18* | AT1G48350 | CCTCTGGACCAACCATTGAG | GTGATACCTTTCTCCAAGCAAGA |
| *RPL28* | AT2G33450 | CCGTATCTCCCTTCCTTCGT | AACTTTGTTTGCTCTGTTTGCTT |
| *RPS6* | AT5G46470 | GTGACTTGAATGAAGAAAGGATGA | GGTAATCTTGTACTTTCTGGTTAATGC |
| *TIC 110* | AT1G06950 | GTTGAACTCTGCTGTACCGGA | CTTTCGTCACAGAGGCAGGAT |
| *TOC159* | AT4G02510 | GAGGCCTCAAGAACCATTGGA | ACAACAAGTAAGGGAGAGGCG |
| *CPN10* | AT2G44650 | GCTTCCACTTTCGTCTGCTCT | GGTTAGCCGTCGAAGGAGTAG |
| *PIF4* | AT2G43010 | AGATACAGAGCCCGGTACAGT | GTTTTGTACCGGGTTTTGGCA |
| *PIF1* | AT2G20180 | CTTGGCTTCATTATCCTCTCC | AGCCTCGAGAAATTCATGAAG |
| *GLK1* | AT2G20570 | ACATCCCACGGTTGATCAGTC | GGATTAGGCATGGCGGTAGAA |
| *GLK2* | AT5G44190 | TAGGAGGAGGAGGGAAGAAGC | AGGCCTGAAGTGACCATGATG |
| *HY5* | AT5G11260 | GATGAGGAGATACGGCGAGTG | GCCGATCCAGATTCTCTACCG |
| *CGA1* | AT4G26150 | CTCGCTCAAGTGGATATCTTC | GCAAATCCTAATCACGCAATC |
| *MYBS1* | AT1G49010 | AGACTCTCATTCCTCTGGATC | CCAATCTCCTTTCCCAAACTT |
| UPL7 | AT3G53090 | TTCAAATACTTGCAGCCAACCTT | CAAAGAGAGGTATCACAAGAGACT |
| PP2AA3 | AT1G13320 | TAACGTGGCCAAAATGATGC | GTTCTCCACAACCGCTTGGT |

Table S2: List of primers used for constructs described in this study

| Gene / Construct | Accession Number | Primer Name | Forward Primer (5′→3′) | Reverse Primer (5′→3′) |
| --- | --- | --- | --- | --- |
| CIA2p::CIA2_cia2cil_ | AT5G57180 | CIA2 Promoter | ggctacggtctcgagta**TGCCACGTCAATGGATAAG** | ggctacggtctctgagcgaatggtgaa**GAAGGGACAAGATAGCTC** |
|  |  | Full genomic sequence | ggctacggtctct**GCTCACTCTCTCTCTCTC** | ggctacggtctcgcttttatccttg**TTAAAGATCTTCTTCGCTAATAAGTTTTTGTTCTCTTTGTCCACTTGG** |
|  |  | 3UTR | ggctacggtctcg**AAAGAGCCTAGATTTATCTTATC** | ggctacggtctcttcca**CTCTGACTTGCTTCTTGAG** |
| 35Sp::CIA2_cia2cil_ | AT5G57180 | 35s promoter | ggctacggtctccagta**TGAGACTTTTCAACAAAGG** | ggctacggtctca**TCTCCAAATGAAATGAACTTC** |
|  |  | Full genomic sequence | ggctacggtctcagagaggacg**TTCACCATTCGCTCACTC** | ggctacggtctcccttttatccttg**TTAAAGATCTTCTTCGCTAATAAGTTTTTGTTCTCTTTGTCCACTTGGAGTGC** |
|  |  | 3UTR | ggctacggtctcc**AAAGAGCCTAGATTTATCTTATC** | ggctacggtctcttcca**CTCTGACTTGCTTCTTGAG** |
| CIA2prom::LUC | AT5G57180 | SALK_004037_LP / NOS_Term_R1 | **CTTCACCATTCGCTCACTCTC** | **TTTATGCTTCCGGCTCGTAT** |
| CILprom::LUC | AT4G28590 | SAIL_228_C01_LP / NOS_Term_R1 | **CCTCTCTGTTTTCGCATCAAC** | **TTTATGCTTCCGGCTCGTAT** |

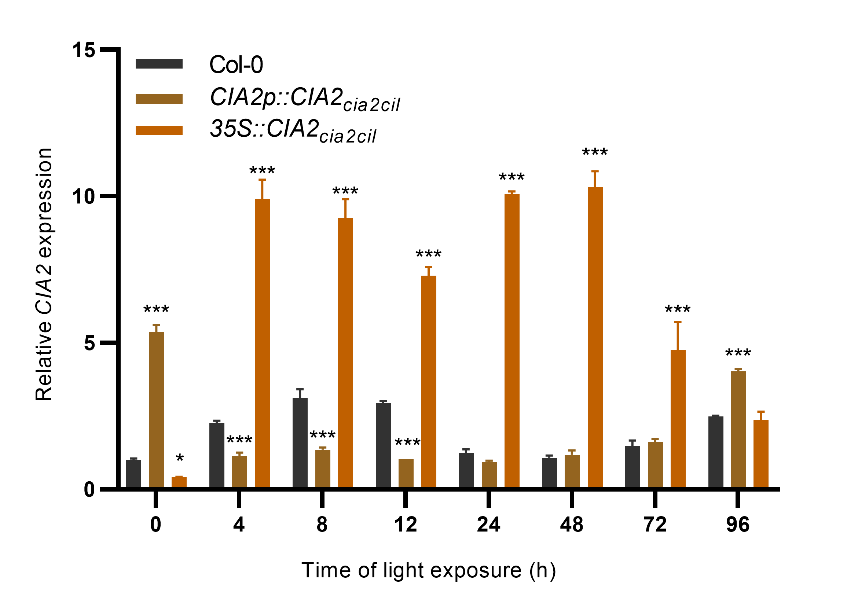

**Figure S1:** Analysis of CIA2 expression in complementation lines (*CIA2p::CIA2_cia2cil_* and *35Sp::CIA2_cia2cil_* ) during the first 96 hours of de-etiolation (T0, T4, T8, T12, T24, T48, T72 and T96). Mean values were derived from 9 measurements (n = 9), and statistical significance (ANOVA and Tukey HSD test) is shown relative to wild-type at each time point analyzed (P < 0.05 (*), P < 0.005 (**), or P < 0.001 (***)).

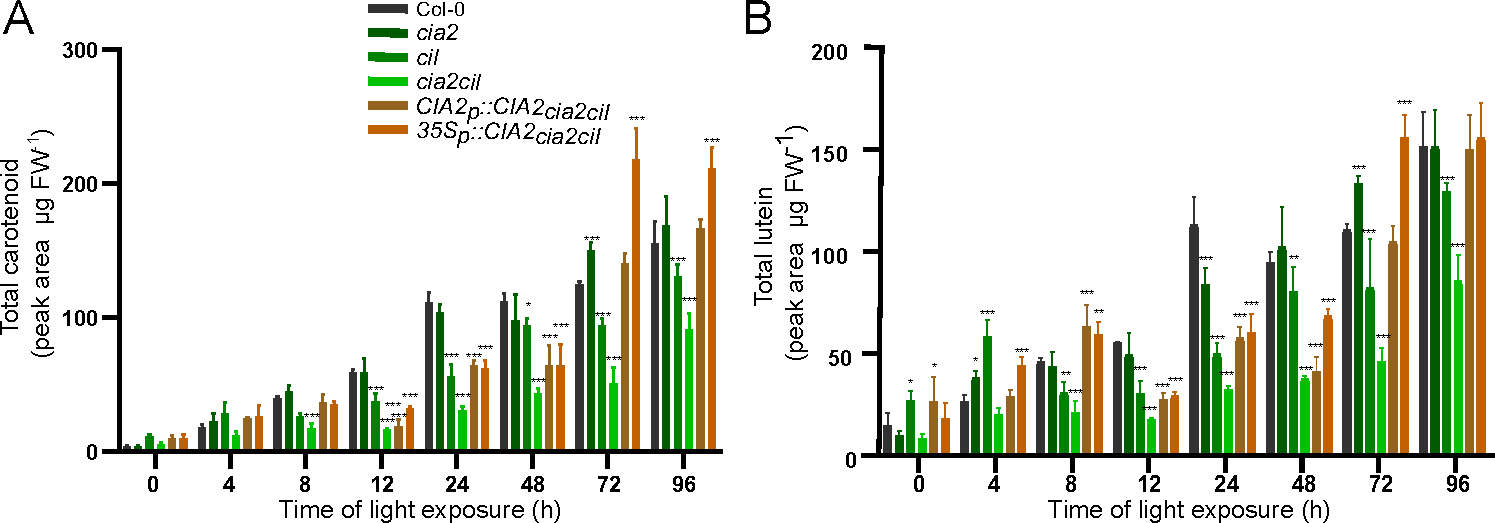

**Figure S2:** Pigment analysis. Content of (A) carotenoids and (B) lutein in Col-0, *cia2*, *cil, cia2cil*, *CIA2p::CIA2_cia2cil_* and *35Sp::CIA2_cia2cil_* seedlings during T0, T4, T8, T12, T24, T48, T72 and T96. Mean values were derived from 9 measurements (n = 9), and statistical significance (ANOVA and Tukey HSD test) is shown relative to wild-type (P < 0.05 (*), P < 0.005 (**), or P < 0.001 (***)).

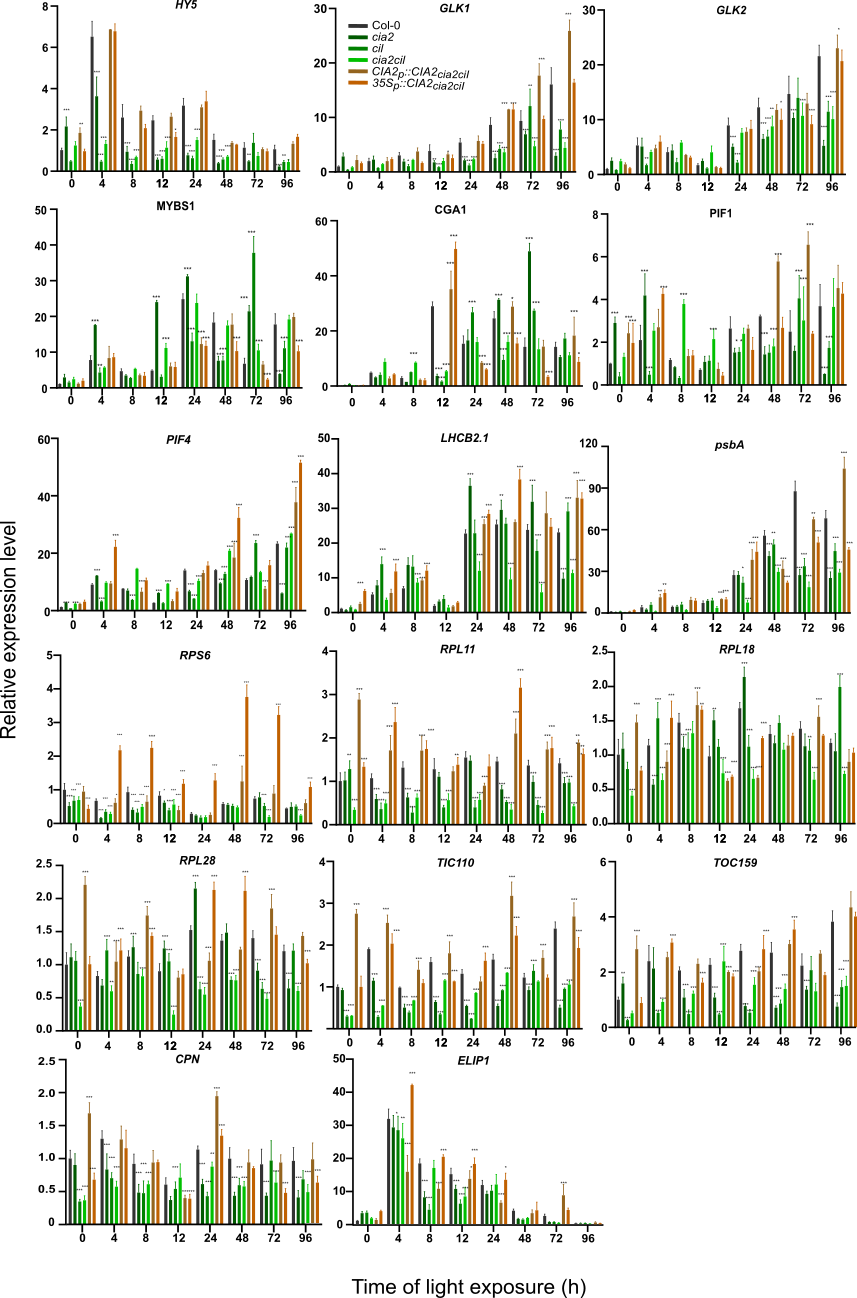

**Figure S3:** Transcriptome Analysis. qRT-PCR analysis of genes expression in the analyzed genotypes in T0, T4, T8, T12, T24, T48, T72 and T96 time points. Mean values were derived from 9 measurements (n = 9), and statistical significance (ANOVA and Tukey HSD test) is shown relative to wild-type at each time point analyzed (P < 0.05 (*), P < 0.005 (**), or P < 0.001 (***)).
